# Single-Cell Transcriptomic Analysis of Livers During NLRP3 Inflammasome Activation Reveals a Novel Immune Niche

**DOI:** 10.1101/2021.03.31.437725

**Authors:** David Calcagno, Angela Chu, Susanne Gaul, Nika Taghdiri, Avinash Toomu, Aleksandra Leszczynska, Benedikt Kaufmann, Alexander Wree, Lukas Geisler, Hal M. Hoffman, Ariel E. Feldstein, Kevin R. King

## Abstract

The NOD-like receptor protein 3 (NLRP3) inflammasome is a central contributor to human acute and chronic liver disease, yet the molecular and cellular mechanisms by which its activation precipitates injury remain incompletely understood. Here, we present single cell transcriptomic profiling of livers from a global transgenic Tamoxifen-inducible constitutively-activated *Nlrp3*^A350V^ mutant mouse, and we investigate the changes in parenchymal and non-parenchymal liver cell gene expression that accompany inflammation and fibrosis. Our results demonstrate that NLRP3 activation causes chronic extramedullary myelopoiesis marked by an increase in proliferating myeloid progenitors that differentiate into neutrophils, monocytes, and monocyte-derived macrophages, results that were corroborated by flow cytometry and histological staining. We observed prominent neutrophil infiltrates with increased Ly6g^HI^ and Ly6g^INT^ cells exhibiting transcriptomic signatures of granulopoiesis typically found in the bone marrow. This was accompanied by a marked increase in Ly6c^HI^ monocytes differentiating into Cd11b^HI^Tim4^HI^Clec4f^HI^ macrophages that express proinflammatory transcriptional programs similar to macrophages of non-alcoholic steatohepatitis (NASH) models. NLRP3 activation also downregulated metabolic pathways in hepatocytes and shifted hepatic stellate cells towards an activated pro-fibrotic state based on expression of collagen and extracellular matrix (ECM) regulatory genes. These results, which highlight abundant neutrophils and extramedullary granulopoiesis define an inflamed and fibrotic hepatic single cell microenvironment, precipitated solely by NLRP3 activation. Clinically, our data support the notion that neutrophils and NLRP3 should be explored as therapeutic targets in NASH-like inflammation.

## Introduction

Non-alcoholic steatohepatitis (NASH) is a highly prevalent inflammatory and fibrotic liver disease that now represents one of the most common reasons for cirrhosis and liver transplantation (1). Despite intense drug development efforts, there are still no FDA approved therapeutics for NASH (2). This is in part due to a lack of consensus regarding which cellular and molecular targets are most pathophysiologically important and actionable. Diverse cell types have been implicated in NASH pathogenesis using bulk methods and surface immunophenotyping; however recently, single cell transcriptomics has emerged to offer a more unbiased characterization of cell-specific transcriptomes in normal and diseased liver (3). Single cell biology is now poised to reveal new secrets about the cellular and molecular mechanisms of liver inflammation and fibrosis (4).

Studies of myeloid cells during inflammation and fibrosis in NASH models have revealed dynamic heterogeneity (5, 6). Kupffer cells, the resident macrophages of the healthy adult liver, decline in number and are replaced by bone marrow-derived monocytes that differentiated into new liver macrophages. Bone marrow-derived liver macrophages have been implicated in exacerbating damage as well as accelerating recovery after pathologic insult (7-9). In the fibrotic liver, single cell analysis also enabled identification and characterization of a “scar-associated” macrophage derived from circulating monocytes with a unique gene signature (10).

NLRP3 activation can serve as a genetic non-diet-induced model of liver inflammation and fibrosis (11). The NLRP3 inflammasome has been implicated in a wide range of inflammatory conditions, such as cardiovascular disease, chronic kidney disease, inflammatory bowel disease, and numerous autoimmune diseases (12-15). Upon activation, the NLRP3 protein binds to the apoptosis-associated speck-like protein containing a C-terminal caspase recruitment domain (ASC) adaptor protein and caspase-1 forming the NLRP3 inflammasome complex, permitting the conversion of pro-interleukin 1β (pro-IL1 β) and pro-interleukin 18 (pro-IL18) into their active forms and inducing cell death in a process termed pyroptosis (16). A subset of humans with the systemic auto-inflammatory disease, cryopyrin associated periodic syndrome (CAPS), carry the Muckle-Wells activating A352V mutation in the NLRP3 protein (17). An analogous mutation was engineered into a mouse model known as *Nlrp3*^A350V^, and its inducible variants are now commonly used as models of NLRP3-induced hyperinflammation (17, 18). In mice, inducible activated *Nlrp3*^A350V^ causes liver fibrosis, pyroptosis, and immune infiltration (11, 19). Small molecule blockade of NLRP3 in mouse models results in reduction of NASH phenotype (20). However, little is known about the diversity of cellular phenotypes within the NLRP3-induced inflamed and fibrosing liver.

Here, we study the inflamed and fibrosed livers of a mouse carrying the global tamoxifen-inducible activated *Nlrp3*^A350V^ mutated gene that was previously demonstrated to exhibit inflammation and fibrosis in the absence of direct liver injury (11). We define single cell transcriptional changes that arise in immune and non-immune cells during NLRP3 inflammasome-activated liver injury. We show that NLRP3 activation results in extramedullary myelopoiesis that supplies an evergreen source of proinflammatory neutrophils and monocyte-derived macrophages leading to hepatocyte metabolic dysfunction and hepatic stellate cell-mediated fibrosis.

## Materials and Methods

### Mouse strains

As previously described, *Nlrp3*^*A350V/+*^CreT knock-in mice (NLRP3-KI) with an alanine 350 to valine (A350V) substitution and an intronic floxed neomycin resistance cassette, in which expression of the mutation does not occur unless the *Nlrp3* mutants are first bred with mice expressing Cre recombinase, were used for this study (21). NLRP3-KI mice were bred to B6.Cg-Tg (Cre/Esr1)5Amc/J mice (obtained from the Jackson Laboratory) to allow for mutant *Nlrp3* expression in adult models after administration of tamoxifen (22).

### Temporal induction of mutant Nlrp3 expression

NLRP3-KI mice and control (*Nlrp3*^*A350V/-*^CreT) mice were injected intraperitoneally with 50 mg/kg tamoxifen-free base (MP Biomedicals) in 90% sunflower seed oil from Helianthus annus (MilliporeSigma) and 10% ethanol daily for 4 days starting at 8 weeks of age as previously described (23).

### Liver sample preparation

NLRP3-KI and control mice were sacrificed 4 weeks after initiation of induction of *Nlrp3* expression. Livers were either prepared for single cell, single nuclei isolation or fluorescence activated cell sorting (FACS) or representative pieces of harvested liver tissue were either (a) fixed in 10% formalin for 24 hours, (b) embedded in OCT on n-Butan nitrogen and then frozen at –80°C, (c) placed in 0.5 mL RNAlater Solution (Invitrogen), or (d) snap-frozen in liquid nitrogen and stored at –80°C. If livers were not prepared for cell isolation, then blood samples (∼0.2 mL) were obtained by heart puncture.

### Liver histology and immunostaining

Livers were sliced in 5-μm sections and were routinely stained for hematoxylin and eosin (H&E). Liver fibrosis was assessed with picosirius red (PSR) staining. For PSR staining, liver sections were incubated for 2 hours at room temperature with an aqueous solution of saturated picric acid containing 0.1% Direct Red (MilliporeSigma). To study liver cell death, terminal deoxynucleotidyl transferase dUTP nick end labeling (TUNEL) assay was performed using manufacturer’s instructions (ApopTag Peroxidase In Situ Apoptosis Detection Kit, MilliporeSigma). Immunohistochemistry (IHC) staining for myeloperoxidase (MPO) (1:200, Thermo Fisher Scientific, catalog RB-373) was performed on formalin-fixed, paraffin-embedded livers according to manufacturer’s instruction and counterstained with Mayer’s Hematoxylin solution (Sigma-Aldrich).

### RNA-seq

Bulk RNA sequencing (RNA-seq) was performed by Q2 Solutions – EA Genomics. Briefly, total RNA was isolated from fresh frozen tissue using the Qiagen miRNeasy Mini Kit. All samples had >100 ng of input RNA and an RNA integrity number (RIN) value ≥ 7.0. Sequencing libraries were created using the IlluminaTruSeq Stranded mRNA method, which preferentially selects for messenger RNA by taking advantage of the polyadenylated tail. Libraries were sequenced using the Illumina sequencing-by-synthesis platform, with a sequencing protocol of 50 bp paired-end sequencing and total read depth of 30 M reads per sample.

RNA-seq data generated in this study were deposited in the Gene Expression Omnibus (http://www.ncbi.nlm.nih.gov/geo) with accession number GSE140742.

### Tissue Processing

Liver cell isolation has been described elsewhere (24). Briefly, mice were anesthetized by ketamine/xylazine injection and perfused *in situ* through the inferior vena cava with sequential Pronase E (0.4 mg/mL, MilliporeSigma) and Collagenase D (0.8 mg/mL, MilliporeSigma) solutions. The entire liver was removed and digested in vitro with Collagenase D (0.5 mg/mL), Pronase E (0.5 mg/mL) and DNAse I (0.02 mg/mL, MilliporeSigma). After 20 minutes, tissue was filtered through a 70-μm mesh. The cell solution was concentrated in preparation for flow cytometry using centrifugation.

Alternatively, mice were intubated and ventilated with 2% isoflurane. After exposing the heart via thoracotomy, livers were perfused with 10mL of ice-cold phosphate-buffered saline (PBS) via a cardiac puncture of the left ventricle. Perfused livers were excised and enzymatically digested in 250 mg aliquots for 1 hour at 37°C under continuous agitation (1200 rpm) in 450 U/ml collagenase I, 125 U/ml collagenase XI, 60 U/ml DNase I, and 60 U/ml hyaluronidase (Sigma), and filtered through a 40 µm nylon mesh in FACS buffer (PBS with 2.5% bovine serum albumin).

Bone marrow cells were collected by flushing femurs with ice-cold PBS. The resulting solution was filtered through a 40 µm nylon mesh and treated with red blood cell (RBC) lysis (BioLegend). Blood was collected by cardiac puncture. The cellular fraction was collected into EDTA-containing tubes (Sigma), and erythrocytes were eliminated using RBC lysis.

### Nuclei Isolation

Mice livers were weighed and minced before flash freezing with liquid nitrogen. Minced samples were resuspended in 0.5 ml nuclei lysis buffer (Millipore Sigma, Nuclei EZ prep, NUC101), 0.2 U/µl RNAse inhibitor (stock 40U/µl, Enzymatics Y9240L) and homogenized with a 2 ml dounce grinder for 5 strokes with A motor and at least 20 strokes with B motor (Sigma-Aldrich D8938). The lysates were filtered through 100 µm and 50 µm cell strainers (CellTrics filters 04-004-2328, 04-004-2327) after resuspending with another 1 mL of nuclei lysis buffer and 10 minutes of incubation. Then they were centrifuged at 5 00 x g for 5 min at 4°C to pellet nuclei. The nuclear pellet was subsequently resuspended in 1.5 mL nuclei lysis buffer with an incubation of 10 minutes followed by centrifugation at 500 x g for 5 min at 4°C to pellet nuclei. The pellet was subsequently washed once in 0.5 mL of nuclei wash buffer (freshly 2% bovine serum albumin (BSA) in 1xPBS, 0.2 U/µl RNAse inhibitor and 1mM of EDTA) and incubated for 5 minutes without disturbing the pellet. After incubation, 1 mL of nuclei wash buffer and resuspension buffer were added and nuclei were resuspended. The washed nuclei were centrifuged at 500 x g for 5 min at 4°C and resuspended in 1.5 mL nuclei wash buffer. The isolated nuclei were centrifuged at 500 x g for 5 min at 4°C and respectively resuspended with Nuclei wash buffer stained with 10 µg/ml 4′,6-diamidino-2-phenylindole (DAPI) before FACS (5mg/ml, Invitrogen D1306). After sorting using purity mode, DAPI+ nuclei were pelleted at 1000 x g for 15 min at 4°C, resuspended in 2% BSA and trypan blue stained nuclei suspension were quality controlled and counted using hemocytometer (Hausser Scientific 3110V).

### Flow Cytometry

For single cell/nuclei barcoding, cellular or nuclei suspensions were stained with DAPI. We enriched live, single cells/nuclei by sorting FSC-W^LO^, DAPI^LO^ cells or DAPI^HI^ nuclei using a Sony MA900. For FACS analysis of leukocyte subsets, cellular suspensions were stained at 4°C in the dark in FACS buffer (PBS with 2.5% bovine serum albumin) with DAPI to exclude dead cells, Ter119 (BioLegend, clone TER-119) to remove unlysed red blood cells, and mouse hematopoietic lineage markers directed against B220 (BioLegend, clone RA3-6B2), Cd49b (BioLegend, clone DX5), Cd90.2 (BioLegend, clone 53-2.1), NK1.1 (BioLegend, clone PK136). Secondary staining of leukocyte subsets was performed using Cd11b (BioLegend, clone M1/70), Ly6g (BioLegend, clone 1A8), F4/80 (Biolegend, clone BM8), Ly6c (BioLegend, clone HK1.4 or BD Bioscience, clone AL-21), and Tim4 (Biolegend, clone RMT4-54).

FACS analysis of hematopoietic progenitors was performed as previously described (24952646). Briefly, cellular suspensions isolated from bone marrow were stained at 4°C in the dark in FACS buffer with lineage markers including phycoerythrin (PE) anti–mouse antibodies directed against directed against B220 (BioLegend, clone RA3–6B2), Cd11b (BioLegend, clone M1/70), Cd11c (BioLegend, clone N418), NK1.1 (BioLegend, clone PK136), TER–119 (BioLegend, clone TER–119), Gr–1 (BioLegend, clone RB6–8C5), Cd8a (BioLegend, clone 53– 6.7), Cd4 (BioLegend, clone GK1.5) and Il7rα (BioLegend, clone A7R34). Then cells were stained with antibodies directed against Ckit (BioLegend, clone 2B8), Sca1 (BioLegend, clone D7), SLAM markers Cd48 (BioLegend, clone HM48–1) and Cd150 (BioLegend, clone TC15– 12F12.2), Cd34 (BioLegend, clone RAM34), Cd16/32 (BioLegend, clone 2.4G2).

### Singe cell and single nuclei sequencing

Single cell/nuclei RNA sequencing (scRNA-seq or snRNA-seq) was performed via microfluidic droplet-based encapsulation, barcoding, and library preparation (10X Genomics). Paired-end sequencing was performed on an Illumina NovaSeq instrument. Low level analysis, including demultiplexing, mapping to a reference transcriptome and eliminating redundant unique molecular identifiers (UMIs), was conducted with 10X CellRanger pipeline. All subsequent scRNA-seq analyses were conducted using the Seurat R package (v3.1) as detailed below.

Total transcript count for each cell was scaled to 10,000 molecules, and ribosomal reads were removed. Raw counts for each gene were normalized to cell-specific transcript count and log-transformed. Cells with between 200 and 4,000 unique genes and <5% mitochondrial counts were retained for further analysis. Highly variable genes were identified with the FindVariableFeatures method by selecting 4,000 genes with the greatest feature variance after variance-stabilizing transformation. Reference-based integration of control and NLRP3-KI datasets utilizing canonical correlation analysis was performed. After scaling and centering expression values for each variable gene, dimensional reduction was performed on integrated data using principal component analysis and cells were clustered according to the Seurat standard workflow. Differentially expressed genes (DEGs) between clusters were determined using a Wilcoxon Rank Sum test. Uniform Manifold Approximation and Projection (UMAP) was used to visualize data in 2D space.

Subclustering was performed by subsetting particular cell-type barcodes, identifying a new set of DEGs within the subset, rescaling their counts, and re-clustering the subset based on newly determined DEGs. In certain instances, dimensional reduction and UMAP was performed again to reveal local structure not captured at higher level dimensional reduction.

For Spearman’s rank correlation analysis, expression of all variable genes was averaged in each subset being compared and ranked by subset-defining DEGs to create a comparator ranked gene list. A Spearman’s rank correlation coefficient (ρ) for each cell was calculated (where d is the distance between gene ranks, n is number of genes) and averaged within subsets.

Collagen formation scores were defined as the summed expression of collagen 1a1, 1a2, 3a1, and 5a2 (*Col1a1, Col1a2, Col3a1*, and *Col5a2)*, normalized by total RNA and multiplied by a scale factor of 10^3^. Cell cycle scores were defined using CellCycleScoring function.

Gene set enrichment analyses (GSEA) were performed using g:Profiler (31066453).

### Statistics

Statistical analysis was performed using GraphPad Prism software (Version 9). All data are represented as mean values +/- standard deviation unless indicated otherwise. A statistical method was not used to predetermine sample size. For comparisons FACS populations, a 2-tailed t-test was used to determine statistical significance. For comparison of single cell derived scores, we implemented a Wilcoxon Rank Sums Test to determine statistical significance. P values less than 0.05 were considered significant and are indicated by asterisks as follows: *p<0.05, **p<0.01, ***p<0.001, ****p<0.0001.

## Results

### NLRP3 activation leads to hepatic fibrosis and cell death

NLRP3-KI and control mice (∼8 weeks of age) were injected with tamoxifen daily for 4 days and then sacrificed 4 weeks later after collecting livers, blood, and bone marrow (**Figure 1A**). As previously demonstrated by our group, severe hepatic fibrosis and hepatic inflammation ensues with a global inducible mutation of *Nlrp3* (19, 25). IHC staining with H&E shows a large immune cell response and liver architectural distortions in the NLRP3-KI model compared to control (**Figure 1B**). PSR staining shows elevated fibrosis throughout the liver in the NLRP3-KI liver (1.8% of area) compared to control (0.3% of area, *p* = < .00001) (**Figure 1C**). TUNEL staining shows a high degree of apoptosis in the NLRP3-KI liver (310 ± 170 apoptotic bodies/field of view (FOV)) compared to the control liver (64 ± 29 apoptotic bodies/FOV, *p* = < .0001) (**Figure 1D**).

**Figure 1:**
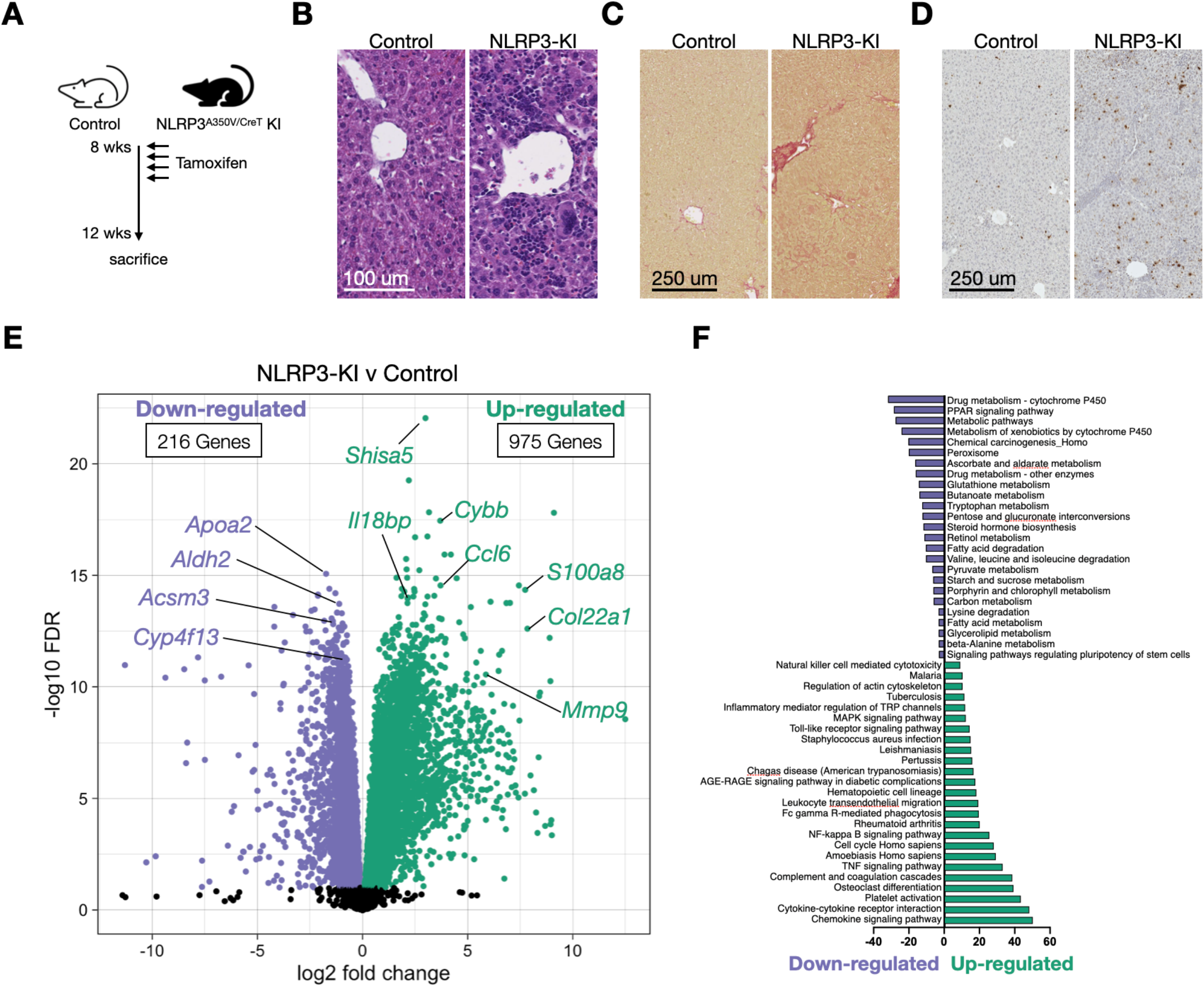
Global activation of NLRP3 inflammasome leads to hepatic inflammation, fibrosis, and cell death. (**A**) Experimental design. (**B-D**) H&E, PSR, and TUNEL stainings of mouse liver sections, respectively. (**E**) Volcano plot of up- and downregulated genes in livers of NLRP3-KI compared to control mice. (**F**) GO analysis of up- (green) and downregulated (purple) genes.

### Bulk RNA-seq implicates innate immune system

To explore the mechanisms underlying NLRP3 inflammatory progression, we performed RNA-seq of the livers from NLRP3-KI and control mice. We detected 975 and 216 genes that exhibited more than 2-fold increase or decrease in mRNA expression, respectively (**Figure 1E** and Table S1). Gene ontology (GO) analysis revealed increased expression of genes associated with several innate immune pathways including chemokine and cytokine signaling, transendothelial leukocyte migration, tumor necrosis factor (TNF) signaling, nuclear factor Kappa B (NF-κB) signaling, phagocytosis, and complement cascades (**Figure 1F**, green). Meanwhile, genes associated with drug metabolism related to cytochrome P450, glutathione metabolism, amino acid metabolism, steroid synthesis, fatty acid degradation and retinol metabolism were downregulated.

### Parenchymal and non-parenchymal cells of the NLRP3-activated liver

To define the specific cell types contributing to NLRP3-mediated inflammation and its sequelae, we performed RNA sequencing on FACS sorted whole single cells and single nuclei collected from the livers of NLRP3-KI and control mice (**Figure 2A**). We integrated the resulting count matrices to produce a harmonized object composed of control and NLRP3-KI samples (**Figure 2B**). This procedure facilitated downstream analyses by ensuring that comparisons were drawn between like cell types. In total, we obtained 3,297 single cell transcriptomes (1751 Control; 1546 NLRP3-KI; n = 1) and 20,119 single nuclei transcriptomes (12,941 control; 7,178 NLRP3-KI; n = 2). The single cell data was composed predominantly of myeloid cells including neutrophils, monocytes, monocyte-derived macrophages (MdMs), and Kupffer cells (KCs). We also captured lymphocytes (B cells, T cells, NK cells), fibroblasts, and endothelial cells to a lesser extent. Clusters were annotated based on the expression of canonical markers revealed by unbiased clustering and DEG analysis (**Figure 2C**). Within the nuclei data, we similarly observed clusters expressing marker genes associated with endothelial cells and immune cells though the bulk of the data was represented by hepatocytes, hepatic stellates cells (HSCs), and cholangiocytes (**Figure 2D, E**). Because we implemented similar isolation protocols, we reasoned that relative abundance of cell types as measured by single cell barcoding was a valid comparison, though limited by cell type frequency and capture protocol. By quantifying cell cluster membership normalized by total cells per sample, we observed a marked increase in immune cells in the NLRP3-KI compared to control mice (**Figure 2, H**). Of note, neutrophils represented roughly half of the cells captured in the single cell NLRP3-KI sample, a 3-fold increase in relative abundance compared to control. Previous reports have indicated that neutrophils play a major role in NLRP3 inflammasome activated liver injury (23). Therefore, we focused our attention on neutrophils.

**Figure 2:**
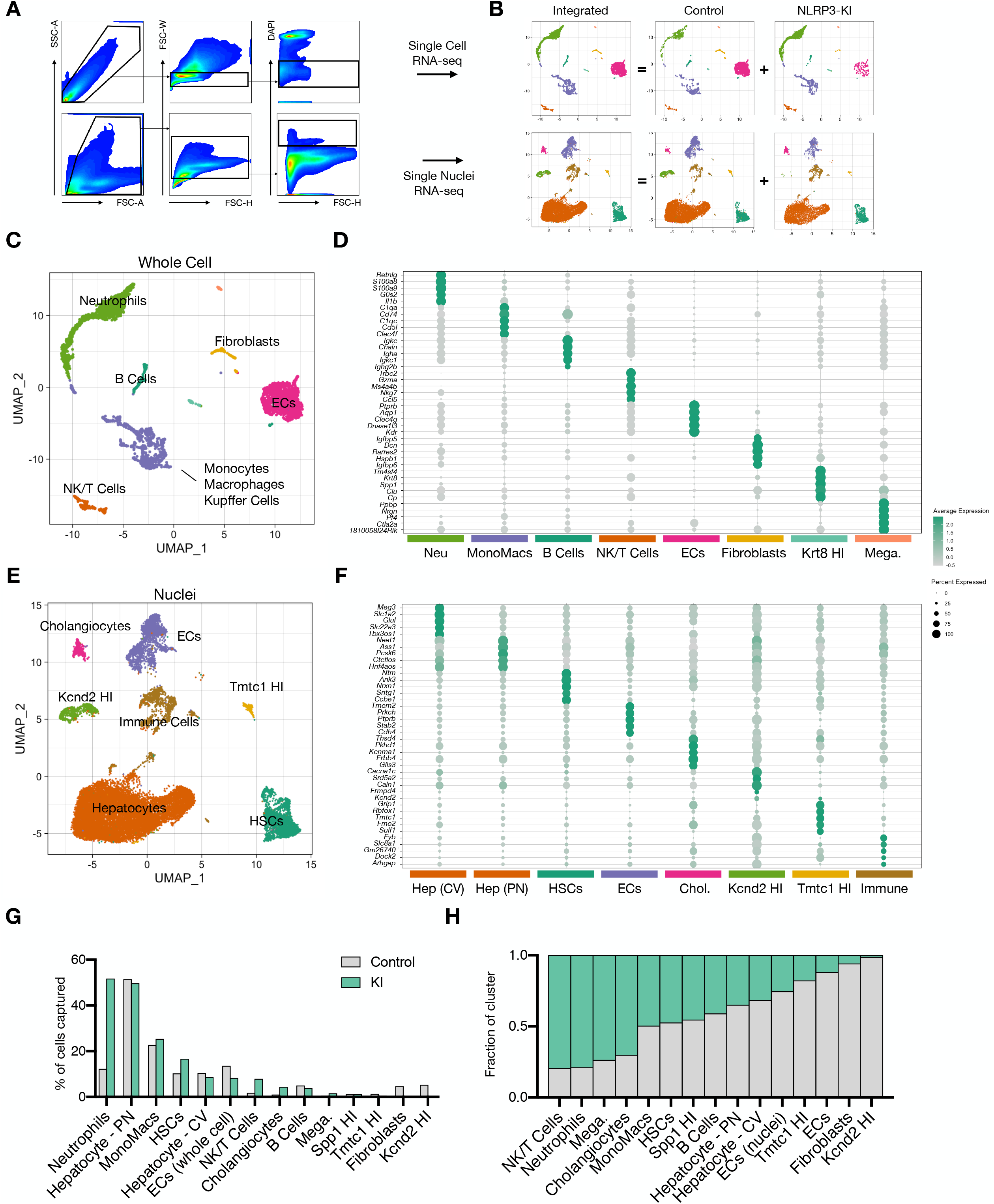
Single cell and nuclei transcriptomics reveals immune, parenchymal, and nonparenchymal compartments in the liver of NLRP3-activated mice. (**A**) Flow cytometry sorting panel for isolation of whole cell (top) and nuclei (bottom) for single cell/nuclei barcoding. (**B**) Integration (batch correction) strategy to classify similar cell types captured in control and NLRP3-activated mice. (**C,E**) UMAP plots of integrated control and NLRP3-activated cells from single cell (**D**) (n = 1; control: 1,751 cells; KI: 1,546 cells) and single nuclei samples (**F**) (n = 1; control: 12,941 nuclei; KI: 7,178 nuclei). (**D,F**) Dot plots displaying top differentially expressed genes in single cell (**D)** and single nuclei (**F**) samples. Data shown is average expression of scaled gene expression by cluster. (**H**) Relative abundance of each cell type, as defined in (**E,G**), as a percentage of total single cell/nuclei transcriptomes captured. (**I**) Fraction of cluster belonging to control and NLRP3-activated (KI) mice. (Neu, neutrophil; ECs, endothelial cells; Mega, megakaryocytes; Hep, hepatocytes; CV, central vein; PN; peripheral node; HSCs, hepatic stellate cells; Chol, cholangiocytes)

### NLRP3 activation causes extramedullary granulopoiesis

Neutrophils are replenished in the bone marrow by a process known as granulopoiesis. In mice, multipotent hematopoietic cells (Lineage^-^Sca1^+^Ckit^+^, LSK) give rise to common myeloid progenitors (CMPs, LS^-^K Cd16/32^+^Cd34^+^) which differentiate into myeloid restricted granulocyte monocyte progenitors (GMPs, LS^-^K Cd16/32^-^Cd34^+^). GMPs then mature into monocytes or neutrophils. Acute liver inflammation typically involves the rapid recruitment of neutrophils, yet the specific functions that neutrophils play during liver injury and the signaling mechanisms for neutrophil recruitment are incompletely understood (26). One study has shown that liver fibrosis was neither worsened nor prevented by diminishing neutrophil recruitment (27), yet other studies have shown increased neutrophil activity leading to more severe hepatic disease (28). To better understand NLRP3-induced neutrophilia, we bioinformatically isolated and reclustered neutrophil transcriptomes defined as *Retnlg*^HI^*Cxcr2*^HI^*Il1β*^HI^*Csf3r*^HI^ cluster (**Figure 3A, 2D**). This resulted in a continuum of neutrophil subtypes, which we divided into 6 clusters and labeled as immature (clusters 1 and 2), intermediate (clusters 3 and 4) and mature (cluster 5 and 6) based on subsequent analyses (**Figure 3B**). We hypothesized that NLRP3 induction prompts extramedullary granulopoiesis. DEG analysis suggested that clusters self-organized onto an axis of maturation similar to granulopoiesis of the bone marrow (**Figure 3C**). To test this, we compared single cell transcriptomes of neutrophils physically isolated from bone marrow (BM) and blood of mice absent of injury, and indeed found a strong quantitative similarity between tissue-specific transcriptomes and the continuum within the liver: clusters 1-3 were most similar to BM neutrophils while cluster 4-6 were most similar to blood neutrophils (**Figure 3D**). Relative to control mice, NLRP3 induction resulted in an increase in replicating immature (*Stmn1*^HI^*Top2a*^HI^*Tubb5*^HI^*Ptma*^HI^*Ppia*^HI^*H2afz*^HI^) and intermediate (*Ly6g*^HI^*Mmp8*^HI^*Retnlg*^HI^) neutrophils (**Figure 3E**). Further, FACS analysis of immune cells isolated from liver of NLRP3-KI and control mice showed a significant increase in Cd11B^HI^ liver leukocytes, as well as Cd11b^HI^Ly6g^HI^ and Ly6g^INT^ neutrophils (**Figure 3F, G**). MPO staining confirmed a large infiltrate of neutrophils in the NLRP3-KI liver (480 ± 320 cells/FOV) compared to control (2 ± 2 cells/FOV, *p* = 0.0465) (**Figure 3H, I**). The quantity of CMPs and GMPs were significantly elevated in the liver of NLRP3-KI mice (**Figure 3J**). The quantity of hematopoietic cells (LSKs) was elevated in NLRP3-KI, though not to a statistically significant degree (p = .0678). To understand the source of progenitor cells, we analyzed leukocytes isolated from peripheral blood. NLRP3 induction increased the number of blood neutrophils (Ly6g^HI^ and Ly6g^INT^) and LS^-^K progenitors (p = .0558, **Figure 3K**). FACS analysis of bone marrow progenitors revealed a roughly 8-fold increase in GMP production of NLRP3-KI mice relative to control (**Figure S1**). From this data, we concluded that NLRP3-activation causes myeloid progenitors to infiltrate the liver, replicate and differentiate into mature neutrophils.

**Figure 3:**
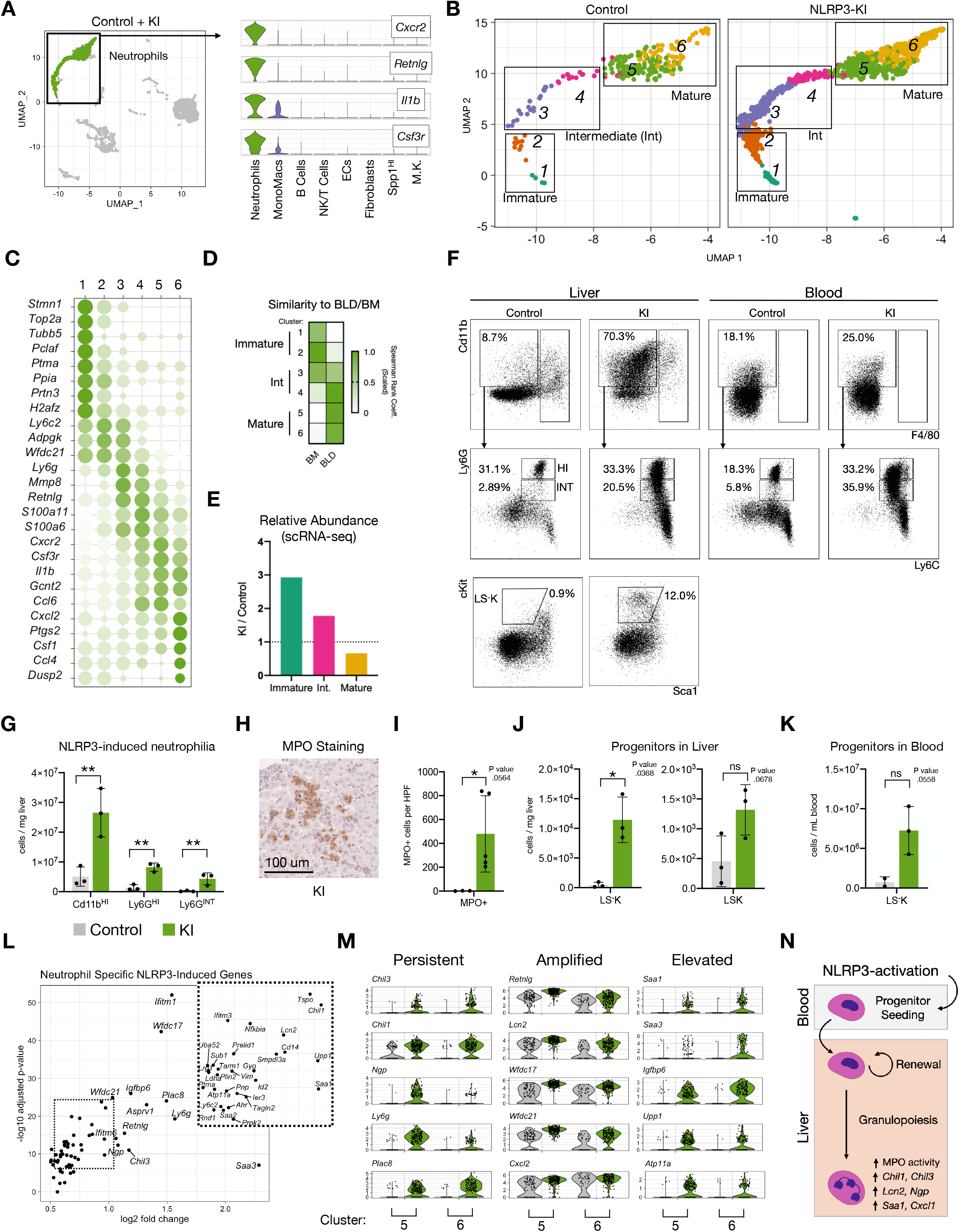
NLRP3 inflammasome activation induces emergency granulopoiesis. (**A**) UMAP and violin plots highlighting neutrophils and neutrophil defining genes for subsequent analyses. (**B**) UMAP plot of subsetted and reclustered neutrophils split by condition. (**C**) Dot plot of DEGs demonstrating continuum of maturation. (**D**) Spearman rank coefficients (row normalized) comparing single cell transcriptomes of neutrophils shown in (B) to neutrophils isolated from bone marrow and peripheral blood of WT mice. (**E**) Relative abundance of immature (clusters 1,2), intermediate (cluster 3,4), and mature (clusters 5,6) in control and NLRP3 induced mice, normalized to control sample. (**F**) FACS analysis of myeloid cells and progenitors of livers and blood isolated from control and KI mice (n=3). (**G**) Quantification of Cd11b^HI^, Ly6g^HI^ and Ly6g^INT^ cells by FACS analysis. (**H**) Representative image from KI mouse showing MPO staining. (**I**) Quantification of MPO staining (manually counted MPO+ cells per 20X FOV). (**J,K**) Quantification of progenitors in liver (J) and blood (K) by FACS analysis. (**L**) Neutrophil-specific NLRP3-induced genes as determined by Wilcoxon Rank Sums test. (**M**) Violin plots of persistent immaturity markers, amplified maturity markers, or entirely up-regulated genes in KI versus control. (**N**) Graphical abstract. * P value < .05, ** P value < .01, unpaired t tests. (Blood, BLD; bone marrow, BM; Wild-type, WT. Myeloperoxidase, MPO.)

Having established that NLRP3 activation results in chronic myelopoiesis, we next asked if its induction perturbed the transcriptional state of mature neutrophils. We compared the transcriptomes of cells belonging to mature neutrophils (clusters 5 and 6) of control and NLRP3-KI mice. Comparisons between other states were restricted due to the lack of immature cells in the control sample. We found up-regulation of roughly 30 genes including those which have been associated with hepatic inflammation and fibrosis (**Figure 3L**). Chitinase-like protein 3 (*Chil3* or *CHI3L1* in humans) has been shown to be expressed in hepatic macrophages of non-alcoholic fatty liver disease (NAFLD) patients (29) with serum levels of YKL-40 elevated in patients with hepatic fibrosis (30). Further, we bucketed up-regulated genes into three groups: persistent, amplified, and elevated (**Figure 3M**). Persistent genes, such as *Chil3*, chitinase-like protein 1 (*Chil1)*, neutrophilic granule protein (*Ngp)*, and placenta associated 8 (*Plac8)* are gene markers for immature and intermediate neutrophils which typically turn off before reaching maturity but were found to be highly upregulated in the mature neutrophils in the NLRP3-KI sample. Amplified genes, such as resistin-like gamma (*Retnlg)*, WAP four-disulfide core domain 17 and 21 (*Wfdc17, Wfdc21*, respectively), are typical gene markers for mature neutrophils, however, were expressed to a greater degree upon NLRP3 activation. Lipocalin-2 (*Lcn2*) was also found to be highly upregulated in the NLRP3-KI model, which has been shown in NASH models to be an important mediator of interactions between neutrophils and hepatic macrophages via CXC chemokine receptor 2 (CXCR2) signaling (31). Elevated genes, such as serum amyloid A1 and A3 (*Saa1, Saa3*, respectively), were notably upregulated compared to control neutrophils. Serum amyloid A (SAA) is an acute phase reactant that can activate the NLRP3 inflammasome cascade, induces synthesis of cytokines, and is a neutrophil chemotactic agent (32). Thus, it is likely that NLRP3 activation has pluripotent effects on granulopoiesis, immune signaling, and neutrophil granule function.

Taken together, this data demonstrates that NLRP3 activation results in hepatic extramedullary granulopoiesis whereby the bone marrow releases myeloid progenitors which seed in the liver, replicate, and differentiate into mature neutrophils with heightened proinflammatory functions (**Figure 3N**).

### NLRP3 activation shifts liver macrophage origins

Given that neutrophils and monocytes share a common progenitor, we next shifted our attention to monocytes, MdMs, and KCs. KCs, the liver’s resident macrophage, derive from monocytes that seed the embryonal liver and are later able to self-replenish. They are phenotypically distinct and have the capacity to express proinflammatory and anti-inflammatory transcriptional programs (33, 34). The balance of proinflammatory KCs and anti-inflammatory KCs helps regulate liver inflammation (35). Recently, several studies have shown that during acute liver injury and NASH, macrophages derived from bone-marrow derived monocytes are critical mediators of pathogenesis (7, 9, 36, 37). These MdMs have been categorized into several phenotypic groups including proinflammatory, wound-healing, and immunosuppressive (38). To examine NLRP3-induced heterogeneity in macrophages, we bioinformatically isolated and reclustered *Cd68*^HI^*C1qa*^HI^ cells as identified during coarse clustering (**Figure 4A, 2D**). These cells spontaneously arranged into 6 clusters which based on DEG analysis we *post hoc* annotated as follows: replicating monocyte progenitors (MPs, *Stmn1*^HI^*Top2a*^HI^), Ly6c^HI^ monocytes (*Ly6c2*^HI^*Ccr2*^HI^), MdM1s (*Ccr2*^HI^*Adgre1*^HI^), MdM2s (*Ccr2*^HI^*Adgre1*^HI^*Cxcl1*^HI^), KCs (*Timd4*^HI^*Clec4f*^HI^), and Ly6c^LO^ cells (*Irf8*^HI^*H2-Eb1*^HI^) (**Figure 4B, C**). The control mice, as expected, were predominately composed of KCs. In contrast, we observed a dramatic shift to MPs, monocytes, and MdMs in NLRP3-KI mice (**Figure 4D**). RNA velocity analysis confirmed the transitory nature of monocytes and MdMs in our dataset (**Figure S2A**). We validated these observations by FACS analysis of monocyte and macrophage subsets. F4/80^LO^Cd11b^HI^Ly6c^HI^ cells were significantly more abundant in NLRP3-activated livers (**Figure 4E, F**). While the quantity of F4/80^HI^ and F4/80^HI^TIM4^HI^ cells were unaltered by NLRP3 activation, the Cd11b mean fluorescent intensity of F4/80^HI^TIM4^HI^ cells increased nearly two fold, suggesting that Tim4^HI^ macrophages were at least partially replenished by Cd11b^HI^ monocytes in concordance with the scRNA-seq data (**Figure 4F,G,H**).

**Figure 4:**
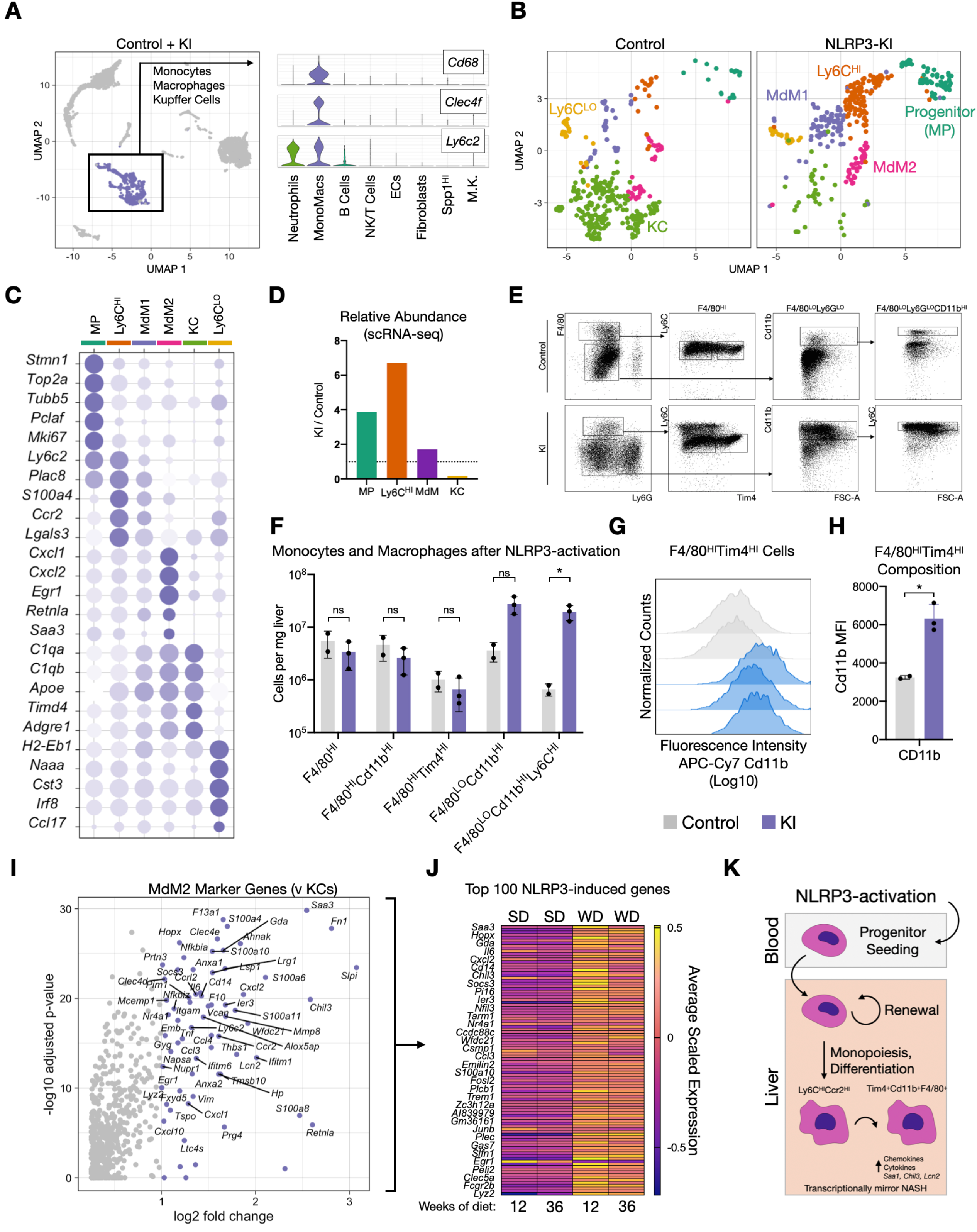
NLRP3 inflammasome activation shifts liver macrophages to monocyte-derived proinflammatory state. (**A**) UMAP and violin plots highlighting monocytes, macrophages, and Kupffer cells and defining genes amongst all captured single cell transcriptomes. (**B**) UMAP plots of subset and reclustered monocyte progenitors (MP), Ly6c^HI^ monocytes, monocyte derived macrophages (MdM) and Kupffer cells (KC), split by condition. (**C**) Dot plot of DEGs showing monocyte differentiation and specialization. (**D**) Relative abundance of each subset normalized to control sample. (**E**) FACS analysis of monocytes, macrophages and KCs in livers of control (grey, n = 2) and KI mice (blue, n=3). (**F**) Quantification of FACS populations. (**G,H**) Mean fluorescence intensity (MFI) of Cd11b (APC-Cy7) in F4/80^HI^TIM4^HI^ cells. (**I**) MdM2-defining markers (Wilcoxon Rank Sums test compared to KC). (**J**) Top 100 MdM2 marker genes applied to monocytes and macrophages from mice treated with standard diet (SD) and Western diet (WD) for 12 and 36 weeks (41). (**K**) Graphical abstract. * P value < .05, ** P value < .01, unpaired t tests.

Recent reports have implicated KC loss and gain of monocyte derived macrophages as a critical mediator of NASH and NAFLD (6, 39). Xion et al characterized the emergence of a triggering receptor expressed on myeloid cells 2+ (*Trem2*+) NASH-associated macrophage (40) while Remmerie et al (41) described an osteopontin-expressing (*Spp1*) MdM with transcriptional similarity to the lipid-associated macrophages (LAM) found in adipose tissue (39) and scar-associated macrophages found in hepatic fibrosis (10). The gene signature of these macrophages, which includes *Spp1, Trem2*, and glycoprotein nonmetastatic melanoma protein B (*Gpnmb)*, are highly expressed upon NLRP3 activation (**Figure S2B,C**). Further, we performed DEG analysis of MdM2 cells compared to KCs and found an up-regulation of over 300 genes including those associated with the monocyte-derived signature (*Ccr2, Ly6c2, Plac8*), prototypical NLRP3-regulated genes (*Il1b, Tnf, Casp4*) and immune recruiting cytokines and chemokines (*Cxcl1, Cxcl2, Ccl2, Ccl3, Ccl4*) (**Figure 4I, S2D**). We applied this MdM2 gene signature to single cell transcriptomes of monocytes and macrophages of mice subjected to standard diet (SD) and Western diet (WD) and found nearly ubiquitous upregulation after 12 and 36 weeks of WD (**Figure 4J**). Taken together, our data demonstrates that NLRP3 activation is sufficient to induce monopoiesis and macrophage specialization similar to those found in models of NASH and hepatic fibrosis (**Figure 4K**).

### Hepatocytes change metabolic function

We next turned to hepatocytes to better understand the effects of NLRP3-driven pathology on the liver itself. Hepatocytes perform myriad functions and have been shown to perform these functions based on spatial location within the liver, such as cholesterol synthesis and urea cycle processing in the highly oxygenated periportal region and bile synthesis and drug metabolism in the chronic hypoxic pericentral region (42). Although liver injury commonly induces regenerative programs, in NASH, hepatocyte-enriched genes are generally downregulated without substantial commensurate proliferation (40, 43). We performed a similar analysis of subsetting and reclustering, as above (**Figure 5A, B**). Subsets anchored to hepatocyte zonation markers as previously described (**Figure S3A**) (42). We performed DEG analysis and found that NLRP3 activation down-regulated ∼2,000 genes while only up-regulating 116 genes, a pattern similar to previous reports in NASH livers (**Figure 5C**) (40). NLRP3-induced hepatocytes displayed dysfunctional metabolism as evidenced by the decrease in expression of genes associated with retinol metabolism, steroid hormone metabolism, glyoxylate metabolism as well as a number of other metabolic pathways (**Figure 5D**). We found little to no evidence of zonation specific regulation (**Figure S3**). A satellite population emerged in the NLRP3-induced sample expressing genes associated with microtubule, tubulin, and cytoskeletal binding, an indication of cellular division and liver regeneration (cluster 7, **Figure 5E**). To test if NASH and NLRP3-induced hepatocytes converged on a similar transcriptional phenotype, we analyzed upregulated and downregulated genes in both models and found no correlation between NLRP3-induced genes and WD (**Figure S4A-C**). This data demonstrates hepatocytes are changing functional priorities from metabolic pathways to cellular regeneration as compensation for apoptosis.

**Figure 5:**
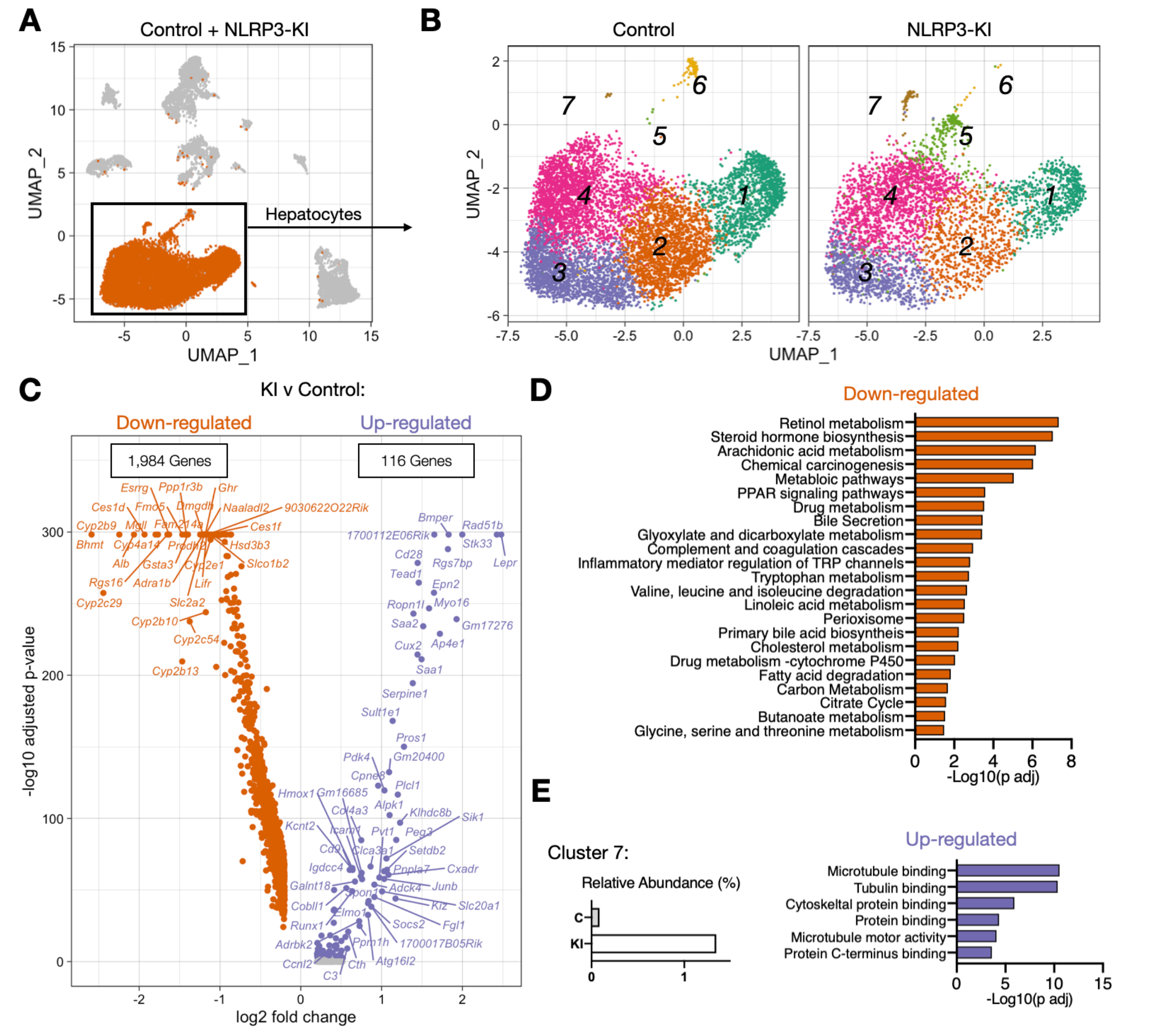
Dysfunctional liver metabolism in the context of chronic NLRP3 inflammasome induction. (**A**) UMAP plot highlighting hepatocytes (snRNA-seq) defined in Figure 2E, F. (**B**) UMAP plots of subset and reclustered hepatocytes, split by condition. Clusters shown represent hepatocyte zonation from central vein to peripheral vein (see figure 3S for more detail). (**C**) Volcano plot of up- and downregulated genes in hepatocytes of NLRP3-KI compared to control mice. (**D**) GSEA of downregulated genes. (**E**) Relative abundance of cluster 7 in control and KI mice (top) and GSEA of cluster defining genes.

### Hepatic stellate cell activation

HSCs have a wide array of functions in the normal liver such as vitamin A storage, vasoregulation, and ECM homeostasis. In response to injury, HSCs specialize into fibrogenic alpha smooth muscle actin (□-SMA)-expressing myofibroblasts (44, 45). This activation is regulated by a complex interplay between HSCs and most cell types found in the liver, and these signaling pathways have been targeted for anti-fibrotic therapies (46). Subsetting and reclustering of HSCs resulted in two broad groups that we annotated as quiescent and activated clusters based on DEG analysis (**Figure 6A,B,C**). Quiescent HSCs expressed high levels of prototypical HSC markers (*Ntm, Ank3, Nrxm1*), while activated HSCs expressed high levels of genes associated with ECM organization (*Gas6, Iga8, Sulf, Smoc2, Eln*) and collagen deposition (*Col1a1, Col1a2, Col3a1, Col5a2*) (**Figure 6C,D**). Some HSCs were activated in the control mice, but we observed a nearly two-fold induction, from 6% to 10.5%, upon NLRP3-activation (**Figure 6E**). As an alternative to cluster membership, we constructed a collagen formation score (see methods). The collagen formation score was significantly higher in HSCs from NLRP3-KI mice compared to control mice, both in quiescent and activated states (**Figure 6F,G**). This was supported by the large amount of L-SMA seen in NLRP3-induced livers **(Figure 6H,I)**.

**Figure 6:**
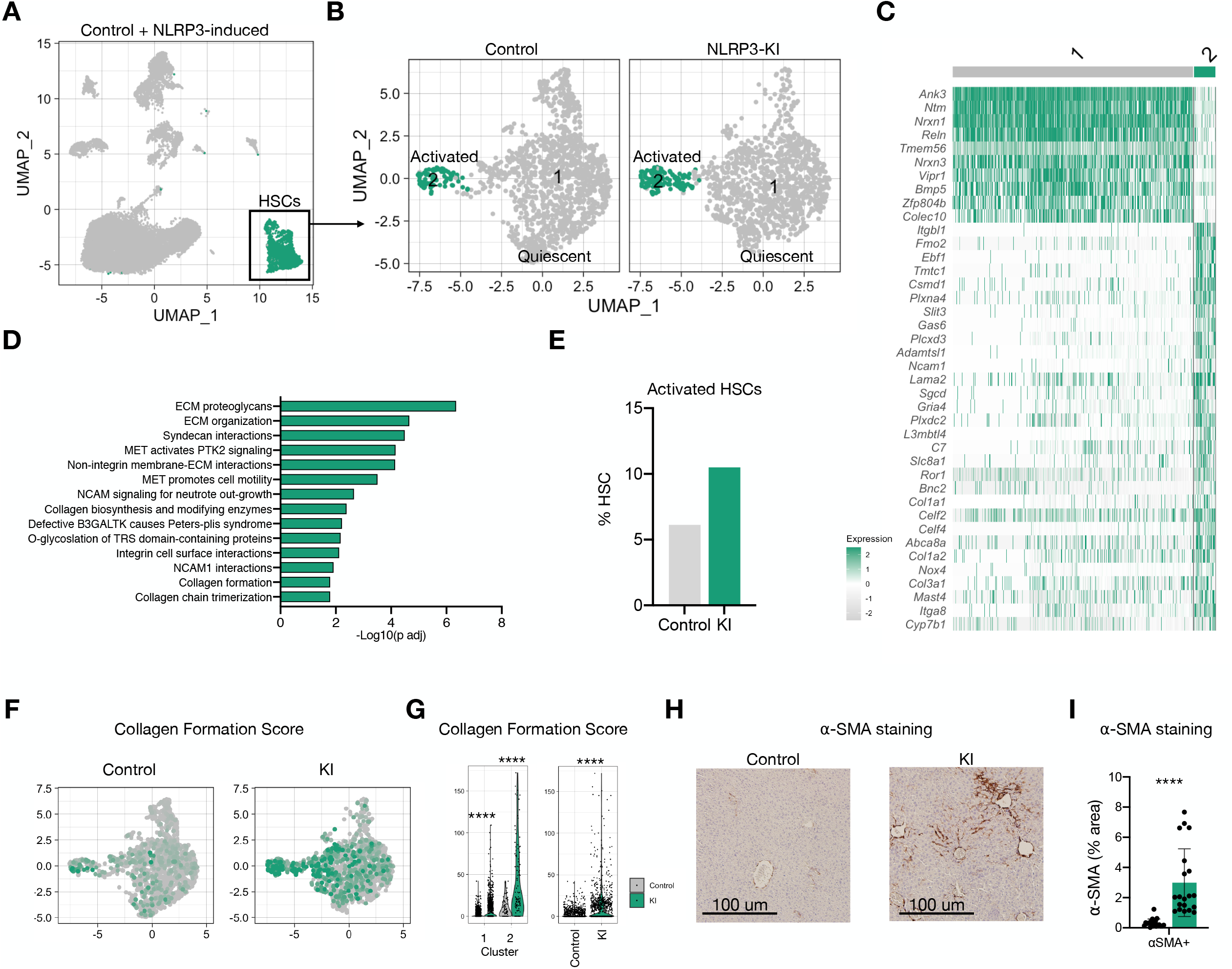
NLRP3 induction activates HSCs and promotes collagen deposition. (**A**) UMAP plots highlighting HSCs for subsequent analyses (snRNA-seq) defined in Figure 2E, F. (**B**) UMAP plots of subset and reclustered HSCs, split by condition. (**C**) Heatmap of DEGs, Wilcoxon Rank Sums Test. (**D**) GSEA of cluster 2 defining genes. (**E**) Relative abundance of activated HSCs (cluster 2). (**F**) Feature plot showing Collagen Formation Score embedded onto UMAP plots, split by condition. (**G**) Violin plots of Collagen Formation Score split by cluster (left) and condition (right). (**H,I**) Representative images of L-SMA staining from control and KI livers. **** P value < .0001, Wilcoxon Rank Sums Test (G) or unpaired t test (I).

## Discussion

Using scRNA-seq, we show that constitutive activation of the NLRP3 inflammasome precipitates chronic extramedullary myelopoiesis, hepatocyte dysfunction, and profibrotic HSC activation in normal liver. This complements a recent report showing that selective inhibition of NLRP3 can reduce liver inflammation and fibrosis in obese diabetic mice (20). Here, we provide further evidence that NLRP3 activation is sufficient to cause NASH-like inflammation and fibrosis even in the absence of steatosis.

scRNA-seq and FACS analyses implicate bone marrow-derived neutrophils and monocytes as key drivers of NLRP3-induced liver pathology. Neutrophils play prominent roles in injury resulting from drug-induced liver injury, physicochemical injury, the response to cell death, and bacterial infection (47-50). They release elastase, MPO, reactive oxygen species (ROS), and secrete cytokines and chemokines, all of which fuel tissue injury, promote inflammation, and recruit additional inflammatory cells (51). Interestingly, there are also reports suggesting that in some contexts, neutrophils play protective roles in resolution of inflammation and in liver repair (52). In the context of NASH, neutrophils have been relatively understudied compared to monocytes and macrophages, in part because neutrophil-preserving assays can be technically challenging. In contrast, monocytes, MdMs and KCs are more mechanically robust cells and are consequently more viable for experimentation. MdMs of NLRP3-activated mice share a remarkably similar transcriptional fingerprint to that of mice fed WD. Future studies are needed to define the relationship of a WD and NLRP3 activation.

Our data provide a number of cell-specific potential driver genes of NLRP3-mediated liver fibrosis. Neutrophils in our NLRP3-activated mice exhibited elevated expression of *Lcn2*, which was recently implicated as a mediator of neutrophil-mediated liver inflammation and macrophage cross-talk in diet-induced NASH (31). Both NLRP3-activated neutrophils and MdMs highly expressed *Chil3*, similar to the human gene *CHI3L1*, which leads to production of chitinase-like protein YKL-40, a biomarker of hepatic fibrosis severity (29, 30). By perturbing chitinase-like proteins in neutrophils and MdMs, future studies can investigate cell-specific contributions and test whether they are essential for NLRP3-mediated liver fibrosis.

Our data raises many additional unanswered questions that will provide inspiration for future experiments. How do NLRP3-activated myeloid cells cause HSC activation and hepatocyte dysfunction? Is NLRP3 essential for this pathogenesis or can other activating mutations drive myeloid infiltration to drive NASH-like inflammation and fibrosis in normal livers? What is clear is that single cell and nuclei transcriptomics will provide an information-rich lens through which to answer these and other mechanistic questions about liver pathobiology.

## Conclusion

Using single cell transcriptomics, we defined and characterized the immune cell subsets within the inflamed liver microenvironment induced by NLRP3 inflammasome activation as well as changes in metabolic status in hepatocytes and HSC phenotypes. These results have important implications for dissecting the mechanisms of liver injury in NLRP3-driven pathologies.

## Supplemental Figures

**Figure S1: FACs analysis of bone marrow progenitors**. (A,B) Representative FACS plots of LSK (Lin^-^Sca1^+^Ckit^+^), LS-K (Lin^-^Sca1^-^Ckit^+^), and subpopulations (GMPs, CMPs, MEPs, Cd48^+^Cd150^-^, Cd48^-^Cd150^+^) from bone marrow (flushed) of control (A) and KI mice (B). (C) Absolute quantification (left) and relative fold change (normalized to control, right) of control (grey) and KI (green) progenitor populations.

**Figure S2: Monopoiesis in NLRP3 activated and similarity to NASH mouse models**. (A) RNA velocity projections embedded on UMAP plots showing progression of monopoiesis. (B) Violin plots of select LAM-associated genes applied to monocytes and macorphages from livers of control (grey) and KI mice (blue, scRNA-seq data). (C) Full LAM signature from Jaitin et al (39). (D) Violin plots of canonical NLRP3 induced genes (left) and cytokines/chemokines (middle, left) of MdMs.

**Figure S3: Effects of NLRP3-induction by hepatocyte zonation**. (A) Heat map showing hepatocyte zonation marker genes from central vein (CV) to peripheral vein (PV). (B) Violin plots quantifying unique molecular indices (UMIs, top) and genes (bottom) split by clusters as defined in Figure 5. (C) Venn diagrams showing up- and down-regulated genes unique and shared across hepatocyte zonations. (C) Volcano plots of up- and downregulated genes (KI v control) by hepatocyte zonation.

**Figure S4: NLRP3-induced genes in hepatocytes and HSCs of NASH model**. Independent analysis of single cell data published in Remmerie et al (41). (A) UMAP plot of coarsely clustered Cd45^-^ cells. (B) Dot plot of cluster defining genes. (C,D) Heatmaps of hepatocyte- (C) and HSC- (D) specific NLRP3 upregulated and downregulated genes applied to correlates in NASH model (left) and diet-induced genes applied to correlates in NLRP3 model (right).

**Figure S5: NLRP3 inflammasome induction recruits and activates T and NK cells**. (A) UMAP plot and violin plots highlighting NK/T cells for subsequent analyses. (B) UMAP plot of integrated (Control + KI) subset and reclustered NK and T cells. (C) Dotplot of DEGs showing NK and T cell specialization. (D) Table quantifying single cell transcriptomes captured in control and NLRP3-induced mice.

## References

1. Friedman SL, Neuschwander-Tetri BA, Rinella M, Sanyal AJ. Mechanisms of NAFLD development and therapeutic strategies. Nat Med 2018;24:908–922.

2. Vuppalanchi R, Noureddin M, Alkhouri N, Sanyal AJ. Therapeutic pipeline in nonalcoholic steatohepatitis. Nat Rev Gastroenterol Hepatol 2021.

3. MacParland SA, Liu JC, Ma XZ, Innes BT, Bartczak AM, Gage BK, Manuel J, et al. Single cell RNA sequencing of human liver reveals distinct intrahepatic macrophage populations. Nat Commun 2018;9:4383.

4. Chu AL, Schilling JD, King KR, Feldstein AE. The Power of Single-Cell Analysis for the Study of Liver Pathobiology. Hepatology 2021;73:437–448.

5. Krenkel O, Hundertmark J, Abdallah AT, Kohlhepp M, Puengel T, Roth T, Branco DPP, et al. Myeloid cells in liver and bone marrow acquire a functionally distinct inflammatory phenotype during obesity-related steatohepatitis. Gut 2020;69:551–563.

6. Seidman JS, Troutman TD, Sakai M, Gola A, Spann NJ, Bennett H, Bruni CM, et al. Niche-Specific Reprogramming of Epigenetic Landscapes Drives Myeloid Cell Diversity in Nonalcoholic Steatohepatitis. Immunity 2020;52:1057–1074 e1057.

7. Tacke F, Zimmermann HW. Macrophage heterogeneity in liver injury and fibrosis. J Hepatol 2014;60:1090–1096.

8. Ju C, Tacke F. Hepatic macrophages in homeostasis and liver diseases: from pathogenesis to novel therapeutic strategies. Cell Mol Immunol 2016;13:316–327.

9. Zigmond E, Samia-Grinberg S, Pasmanik-Chor M, Brazowski E, Shibolet O, Halpern Z, Varol C. Infiltrating monocyte-derived macrophages and resident kupffer cells display different ontogeny and functions in acute liver injury. J Immunol 2014;193:344–353.

10. Ramachandran P, Dobie R, Wilson-Kanamori JR, Dora EF, Henderson BEP, Luu NT, Portman JR, et al. Resolving the fibrotic niche of human liver cirrhosis at single-cell level. Nature 2019;575:512–518.

11. Schuster-Gaul S, Geisler LJ, McGeough MD, Johnson CD, Zagorska A, Li L, Wree A, et al. ASK1 inhibition reduces cell death and hepatic fibrosis in an Nlrp3 mutant liver injury model. JCI Insight 2020;5.

12. Zhou W, Chen C, Chen Z, Liu L, Jiang J, Wu Z, Zhao M, et al. NLRP3: A Novel Mediator in Cardiovascular Disease. J Immunol Res 2018;2018:5702103.

13. Hutton HL, Ooi JD, Holdsworth SR, Kitching AR. The NLRP3 inflammasome in kidney disease and autoimmunity. Nephrology (Carlton) 2016;21:736–744.

14. Zhen Y, Zhang H. NLRP3 Inflammasome and Inflammatory Bowel Disease. Front Immunol 2019;10:276.

15. Shen HH, Yang YX, Meng X, Luo XY, Li XM, Shuai ZW, Ye DQ, et al. NLRP3: A promising therapeutic target for autoimmune diseases. Autoimmun Rev 2018;17:694–702.

16. Schroder K, Tschopp J. The inflammasomes. Cell 2010;140:821–832.

17. Hoffman HM, Mueller JL, Broide DH, Wanderer AA, Kolodner RD. Mutation of a new gene encoding a putative pyrin-like protein causes familial cold autoinflammatory syndrome and Muckle-Wells syndrome. Nat Genet 2001;29:301–305.

18. Hoffman HM, Broderick L. The role of the inflammasome in patients with autoinflammatory diseases. J Allergy Clin Immunol 2016;138:3–14.

19. Wree A, Eguchi A, McGeough MD, Pena CA, Johnson CD, Canbay A, Hoffman HM, et al. NLRP3 inflammasome activation results in hepatocyte pyroptosis, liver inflammation, and fibrosis in mice. Hepatology 2014;59:898–910.

20. Mridha AR, Wree A, Robertson AAB, Yeh MM, Johnson CD, Van Rooyen DM, Haczeyni F, et al. NLRP3 inflammasome blockade reduces liver inflammation and fibrosis in experimental NASH in mice. J Hepatol 2017;66:1037–1046.

21. Brydges SD, Mueller JL, McGeough MD, Pena CA, Misaghi A, Gandhi C, Putnam CD, et al. Inflammasome-mediated disease animal models reveal roles for innate but not adaptive immunity. Immunity 2009;30:875–887.

22. Hayashi S, McMahon AP. Efficient recombination in diverse tissues by a tamoxifen-inducible form of Cre: a tool for temporally regulated gene activation/inactivation in the mouse. Dev Biol 2002;244:305–318.

23. McGeough MD, Pena CA, Mueller JL, Pociask DA, Broderick L, Hoffman HM, Brydges SD. Cutting edge: IL-6 is a marker of inflammation with no direct role in inflammasome-mediated mouse models. J Immunol 2012;189:2707–2711.

24. Mederacke I, Dapito DH, Affo S, Uchinami H, Schwabe RF. High-yield and high-purity isolation of hepatic stellate cells from normal and fibrotic mouse livers. Nat Protoc 2015;10:305–315.

25. Wree A, McGeough MD, Pena CA, Schlattjan M, Li H, Inzaugarat ME, Messer K, et al. NLRP3 inflammasome activation is required for fibrosis development in NAFLD. J Mol Med (Berl) 2014;92:1069–1082.

26. Koyama Y, Brenner DA. Liver inflammation and fibrosis. J Clin Invest 2017;127:55–64.

27. Moles A, Murphy L, Wilson CL, Chakraborty JB, Fox C, Park EJ, Mann J, et al. A TLR2/S100A9/CXCL-2 signaling network is necessary for neutrophil recruitment in acute and chronic liver injury in the mouse. J Hepatol 2014;60:782–791.

28. Zang S, Ma X, Zhuang Z, Liu J, Bian D, Xun Y, Zhang Q, et al. Increased ratio of neutrophil elastase to alpha1-antitrypsin is closely associated with liver inflammation in patients with nonalcoholic steatohepatitis. Clin Exp Pharmacol Physiol 2016;43:13–21.

29. Kumagai E, Mano Y, Yoshio S, Shoji H, Sugiyama M, Korenaga M, Ishida T, et al. Serum YKL-40 as a marker of liver fibrosis in patients with non-alcoholic fatty liver disease. Sci Rep 2016;6:35282.

30. Johansen JS, Christoffersen P, Moller S, Price PA, Henriksen JH, Garbarsch C, Bendtsen F. Serum YKL-40 is increased in patients with hepatic fibrosis. J Hepatol 2000;32:911–920.

31. Ye D, Yang K, Zang S, Lin Z, Chau HT, Wang Y, Zhang J, et al. Lipocalin-2 mediates non-alcoholic steatohepatitis by promoting neutrophil-macrophage crosstalk via the induction of CXCR2. J Hepatol 2016;65:988–997.

32. Eklund KK, Niemi K, Kovanen PT. Immune functions of serum amyloid A. Crit Rev Immunol 2012;32:335–348.

33. Dixon LJ, Barnes M, Tang H, Pritchard MT, Nagy LE. Kupffer cells in the liver. Compr Physiol 2013;3:785–797.

34. Tsutsui H, Nishiguchi S. Importance of Kupffer cells in the development of acute liver injuries in mice. Int J Mol Sci 2014;15:7711–7730.

35. Wan J, Benkdane M, Teixeira-Clerc F, Bonnafous S, Louvet A, Lafdil F, Pecker F, et al. M2 Kupffer cells promote M1 Kupffer cell apoptosis: a protective mechanism against alcoholic and nonalcoholic fatty liver disease. Hepatology 2014;59:130–142.

36. Karlmark KR, Weiskirchen R, Zimmermann HW, Gassler N, Ginhoux F, Weber C, Merad M, et al. Hepatic recruitment of the inflammatory Gr1+ monocyte subset upon liver injury promotes hepatic fibrosis. Hepatology 2009;50:261–274.

37. Seki E, De Minicis S, Gwak GY, Kluwe J, Inokuchi S, Bursill CA, Llovet JM, et al. CCR1 and CCR5 promote hepatic fibrosis in mice. J Clin Invest 2009;119:1858–1870.

38. Mosser DM, Edwards JP. Exploring the full spectrum of macrophage activation. Nat Rev Immunol 2008;8:958–969.

39. Jaitin DA, Adlung L, Thaiss CA, Weiner A, Li B, Descamps H, Lundgren P, et al. Lipid-Associated Macrophages Control Metabolic Homeostasis in a Trem2-Dependent Manner. Cell 2019;178:686–698 e614.

40. Xiong X, Kuang H, Ansari S, Liu T, Gong J, Wang S, Zhao XY, et al. Landscape of Intercellular Crosstalk in Healthy and NASH Liver Revealed by Single-Cell Secretome Gene Analysis. Mol Cell 2019;75:644–660 e645.

41. Remmerie A, Martens L, Thone T, Castoldi A, Seurinck R, Pavie B, Roels J, et al. Osteopontin Expression Identifies a Subset of Recruited Macrophages Distinct from Kupffer Cells in the Fatty Liver. Immunity 2020;53:641–657 e614.

42. Halpern KB, Shenhav R, Matcovitch-Natan O, Toth B, Lemze D, Golan M, Massasa EE, et al. Single-cell spatial reconstruction reveals global division of labour in the mammalian liver. Nature 2017;542:352–356.

43. Ramachandran P, Matchett KP, Dobie R, Wilson-Kanamori JR, Henderson NC. Single-cell technologies in hepatology: new insights into liver biology and disease pathogenesis. Nat Rev Gastroenterol Hepatol 2020;17:457–472.

44. Puche JE, Saiman Y, Friedman SL. Hepatic stellate cells and liver fibrosis. Compr Physiol 2013;3:1473–1492.

45. Tsuchida T, Friedman SL. Mechanisms of hepatic stellate cell activation. Nat Rev Gastroenterol Hepatol 2017;14:397–411.

46. Higashi T, Friedman SL, Hoshida Y. Hepatic stellate cells as key target in liver fibrosis. Adv Drug Deliv Rev 2017;121:27–42.

47. Lammermann T, Afonso PV, Angermann BR, Wang JM, Kastenmuller W, Parent CA, Germain RN. Neutrophil swarms require LTB4 and integrins at sites of cell death in vivo. Nature 2013;498:371–375.

48. Dickson BV, Clack JA, Smithson TR, Pierce SE. Functional adaptive landscapes predict terrestrial capacity at the origin of limbs. Nature 2021;589:242–245.

49. Patel SJ, Milwid JM, King KR, Bohr S, Iracheta-Vellve A, Li M, Vitalo A, et al. Gap junction inhibition prevents drug-induced liver toxicity and fulminant hepatic failure. Nat Biotechnol 2012;30:179–183.

50. Wang J, Hossain M, Thanabalasuriar A, Gunzer M, Meininger C, Kubes P. Visualizing the function and fate of neutrophils in sterile injury and repair. Science 2017;358:111–116.

51. Ley K, Hoffman HM, Kubes P, Cassatella MA, Zychlinsky A, Hedrick CC, Catz SD. Neutrophils: New insights and open questions. Sci Immunol 2018;3.

52. Parthasarathy G, Revelo X, Malhi H. Pathogenesis of Nonalcoholic Steatohepatitis: An Overview. Hepatol Commun 2020;4:478–492.

