## Supplementary figures and images for "Single-Cell Transcriptomic Analysis of Livers During NLRP3 Inflammasome Activation Reveals a Novel Immune Niche"

### Supplemental Figures

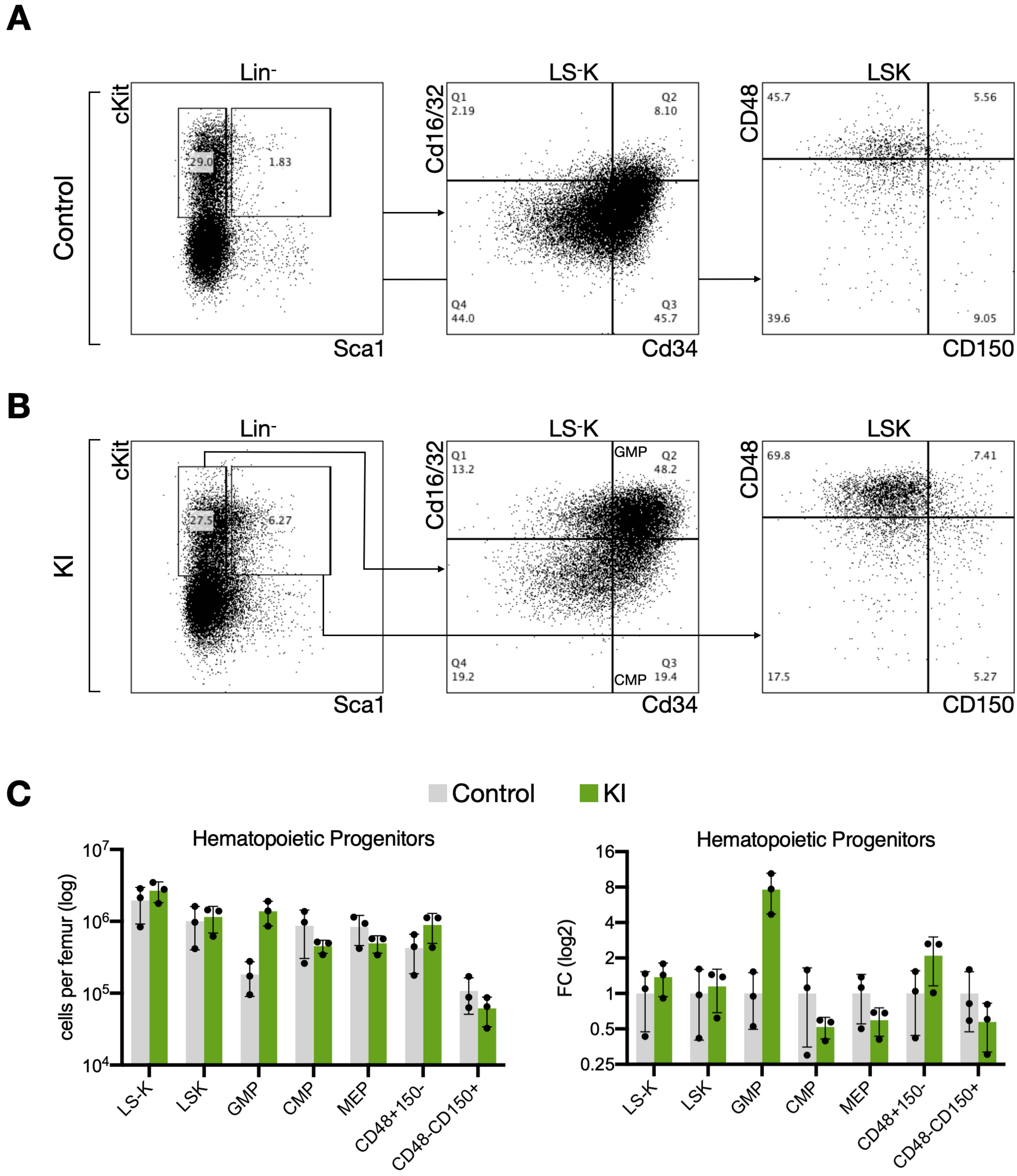

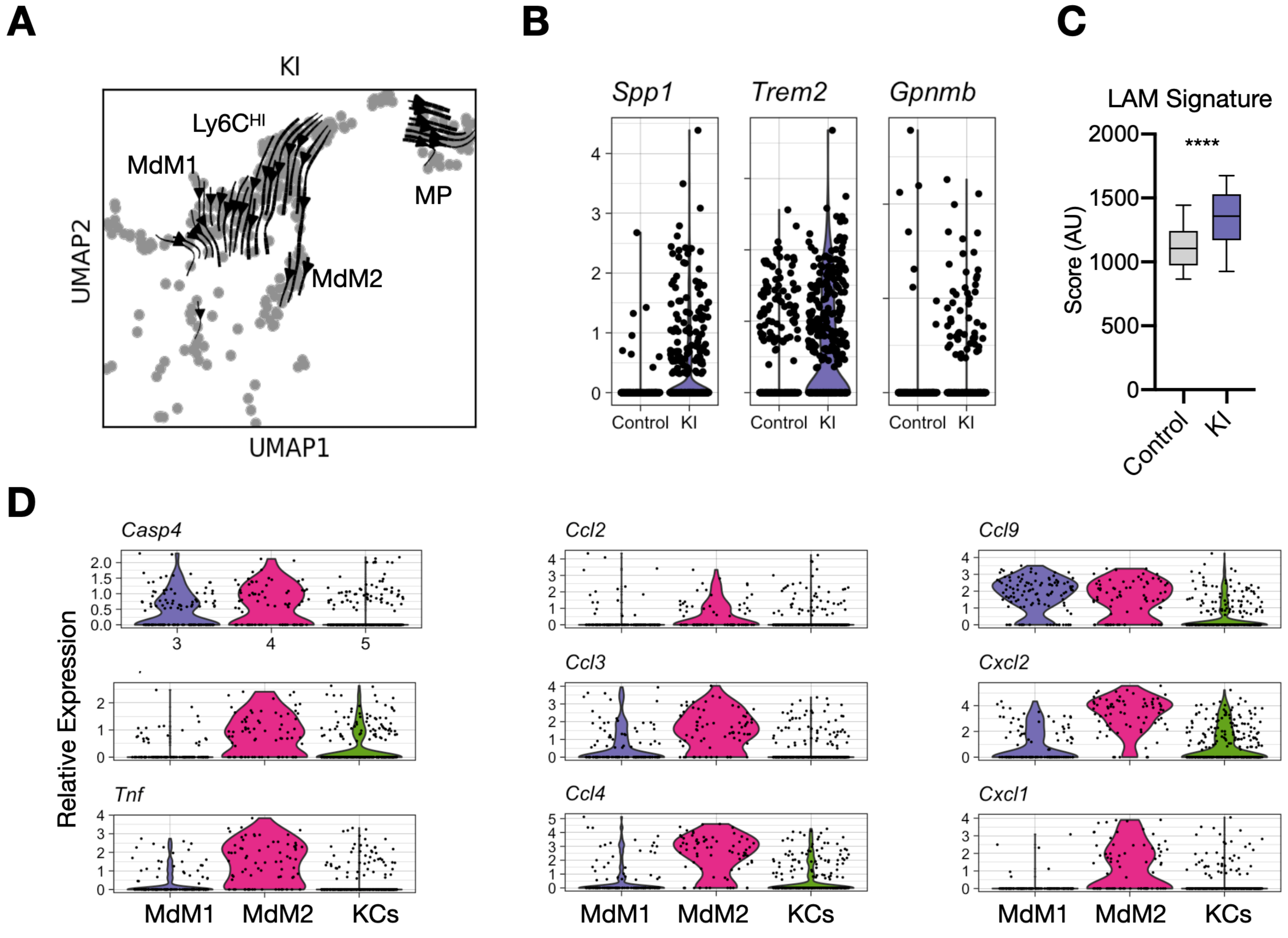

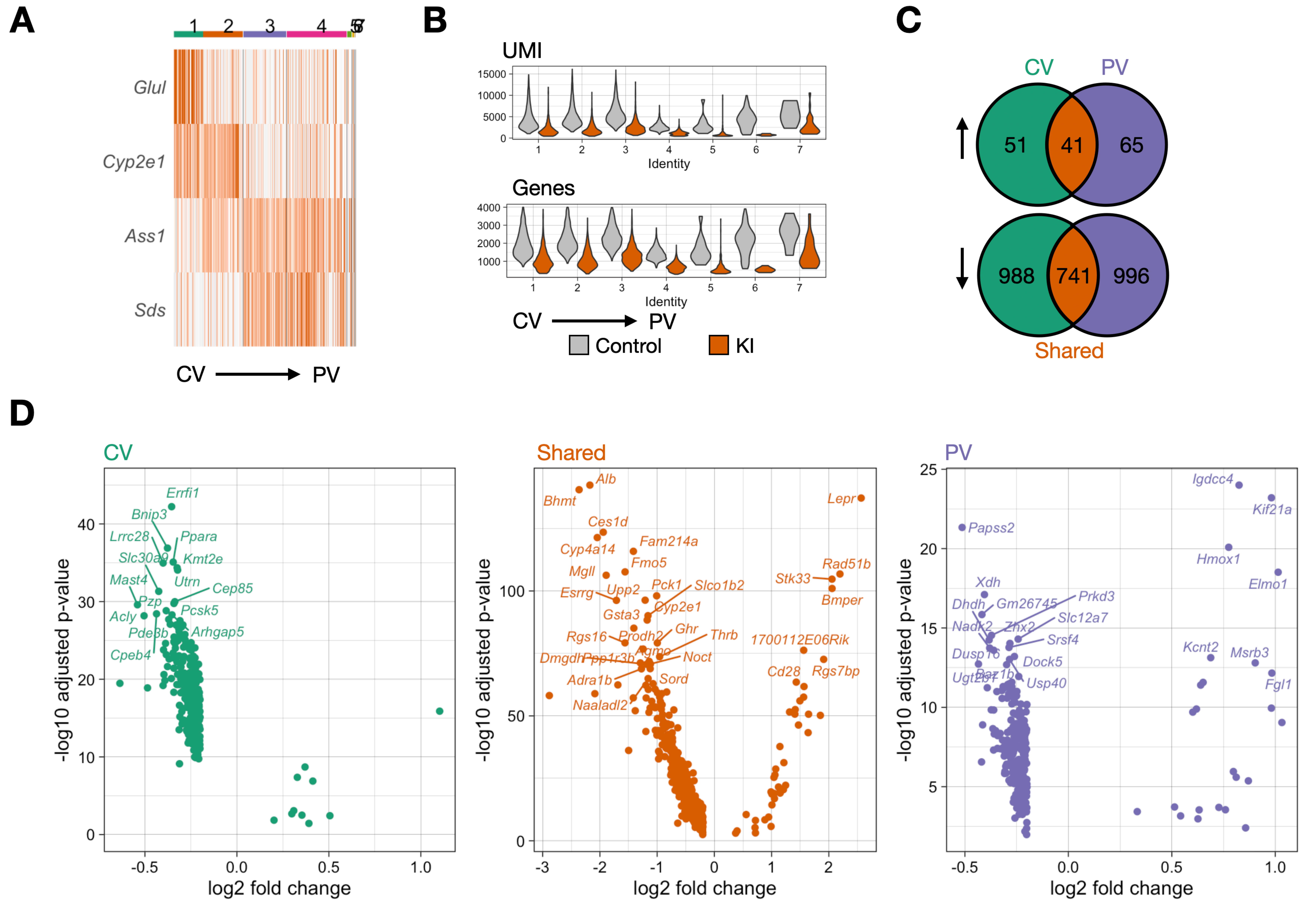

**A**

**B**

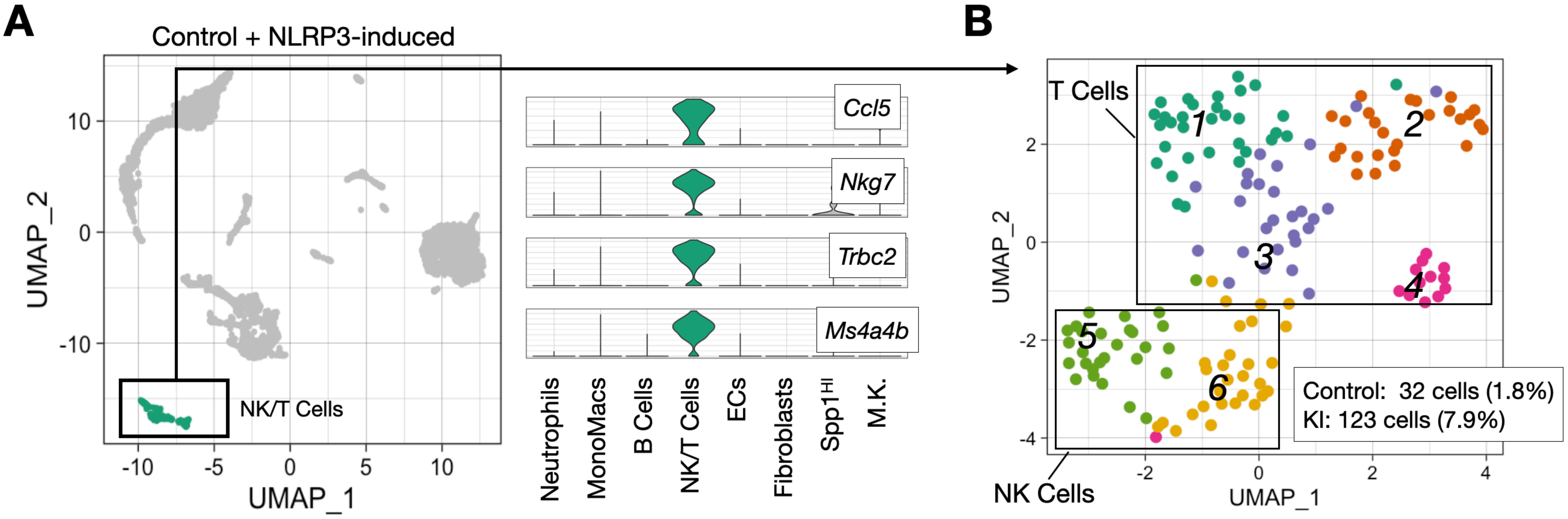


**C**

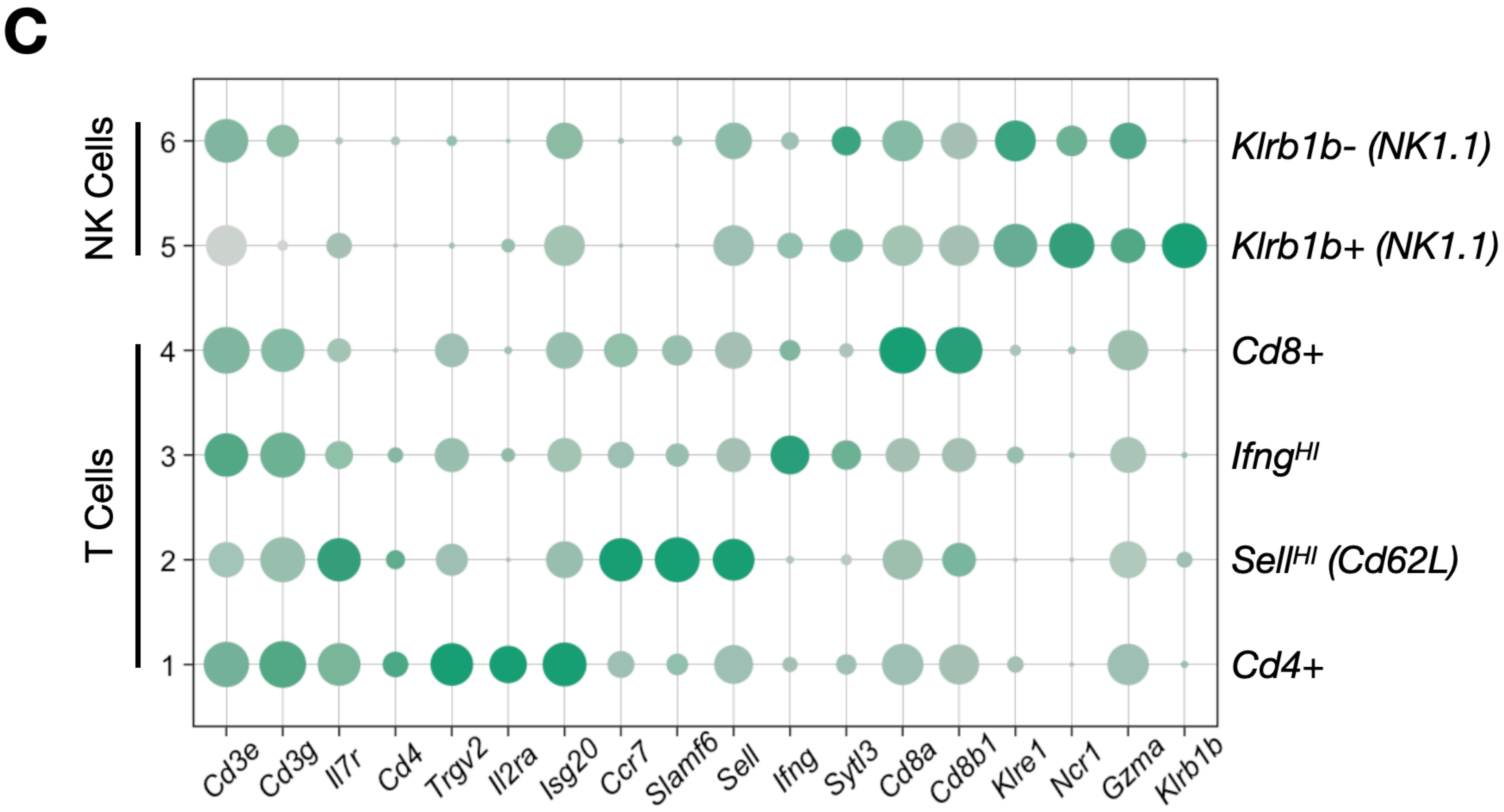


**D**

| | *Cd4+* | *Sell[HI] (Cd62L)* | *Ifng[HI]* | *Cd8+* | *Klrb1b+ (NK1.1)* | *Klrb1b- (NK1.1)* |
|---|---|---|---|---|---|---|
| Control (1.8%) | 5 | 5 | 8 | 3 | 4 | 7 |
| KI (7.9%) | 27 | 21 | 19 | 10 | 24 | 22 |

#### Figure S4
